# Development of rapid and affordable virus-mimicking artificial positive controls

**DOI:** 10.1101/2023.05.31.543123

**Authors:** Shivani Singh, Daisy Stainton, Ioannis E. Tzanetakis

## Abstract

A major bottleneck in the development of detection assays is the availability of positive controls. Their acquisition can be problematic; their maintenance is expensive and without them assays cannot be validated. Herein we present a novel strategy for the development of virus-mimicking positive controls (ViMAPCs). The time between design and application is less than five days, unlike alternatives which normally take several weeks to obtain and implement. The ViMAPCs provide a realistic representation of natural infection unlike alternatives and allow for an effortless recognition of lab-based contamination. The feasibility and adaptability of the strategy was evaluated using several RNA and DNA viruses. ViMAPCs can be used in diagnostics labs but also in monitoring of pathogen outbreaks where rapid response is of utmost importance.

## INTRODUCTION

The use of high throughput sequencing has accelerated the discovery of new organisms and pathogens including viruses (Villamor et al., 2019). Once identified, those viruses often make it to a list of targeted pathogens and affect movement of biological material and goods across borders (Martin et al., 2016). The sheer number and the global distribution of the new discoveries makes acquisition of infected material a herculean task. The importation of infected tissue or nucleic acids is time consuming, requires specialized permits, and in the case of select agents, it is prohibited unless imported by a high containment facility. To address this issue, development of artificial positive controls (APCs) for detection purposes has increased significantly in recent years.

The most commonly used approach for the development of APCs are plasmids/gBlocks™ that incorporate the sequences of interest. Development of DNA-based APCs is straightforward and even though they provide templates that yield amplicons there are challenges. Contamination is difficult to control in lab settings because of their stability and high concentration; jeopardizing testing integrity (Borst et al., 2004; Caasi et al., 2013 and Chomič et al., 2010). Other APCs used include synthetic oligonucleotide templates which are limited to short targets (Whiley et al., 2010), or in-vitro expression of RNA transcripts (Lee at al., 2011; de León et al., 2013; Kodani and Winchell, 2012; and Snow et al., 2009), however, the latter require specialized skills, are often time consuming and add a layer of complexity which most diagnostic labs are not equipped to apply to their daily operations. Yet, all aforementioned methods do not mimic virus infection and the presence of an intense amplicon does not necessarily reflect natural infection, misrepresenting the sensitivity of the assay.

Here we present an approach to develop rapid and affordable positive controls for PCR assays which negates the above-mentioned issues. This approach is a strategic advancement in terms of developing virus-mimicking artificial positive controls (ViMAPCs) irrespective of the virus genome composition or replication mechanism. This technology has the potential to eliminate the aforementioned barriers and allows for the development of realistic positive controls that mimic natural infection. In addition, these ViMAPC are designed to immediately differentiate between natural infection and control; allowing for easy distinction between infection and contamination. Independent of whether the organism was discovered a few kilometers away or on the other side of the globe ViMAPCs can be developed and used in less than a week. The strategy is straightforward and can be implemented readily by any laboratories that use PCR-based diagnostics.

## METHODS

### Material

Material with known virus infections were acquired from the National Clean Germplasm Repository in Corvallis, Oregon or are part of the virus collection of the virology lab of the Department of Entomology and Plant Pathology at University of Arkansas, Fayetteville, United States. *Fragaria, Ficus, Rosa, Rubus, Vaccinium* and *Vitis* spp. were used in the preparation of ViMAPCs (Supplemental table 1) with total nucleic acid extractions (TNAE) performed as described in Poudel et al. (2013).

### Concept

As this is a proof-of-concept study, the primers used in the development of all the ViMAPC incorporate previously published sequences (Supplemental table 1). The template is virus-infected material readily available in the diagnostic laboratory. In order for the ViMAPC to better mimic natural infection, template and target viruses belong to the same genus and when possible, host. The reverse transcription (RT) primers are chimeric (Supplemental table 2) and are comprised of target sequences at the 5’ and template sequences at the 3’ (Fig. 1A) whereas the PCR amplification is conducted using only the target virus primers (Fig. 1B).

**Figure 1.**
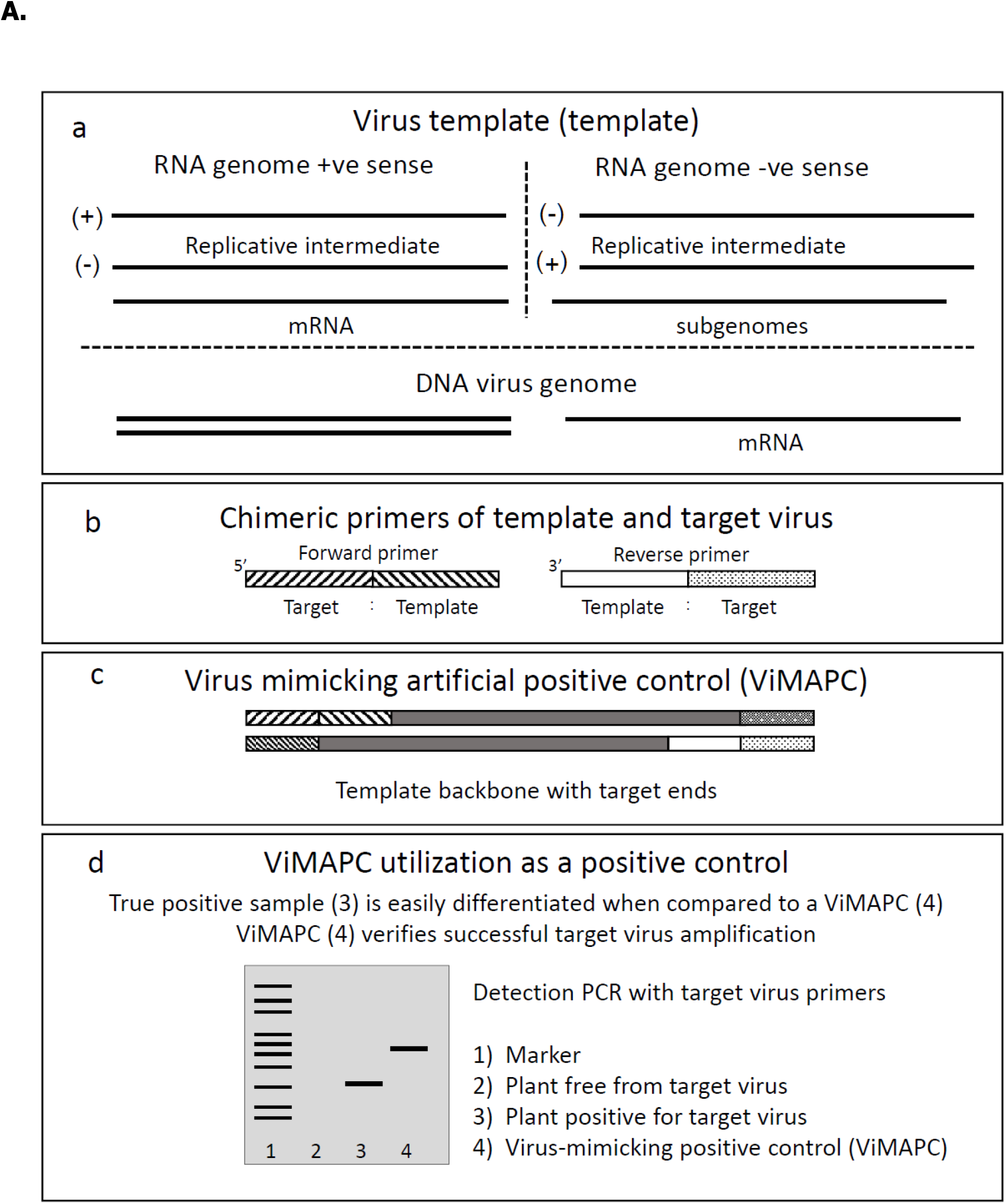
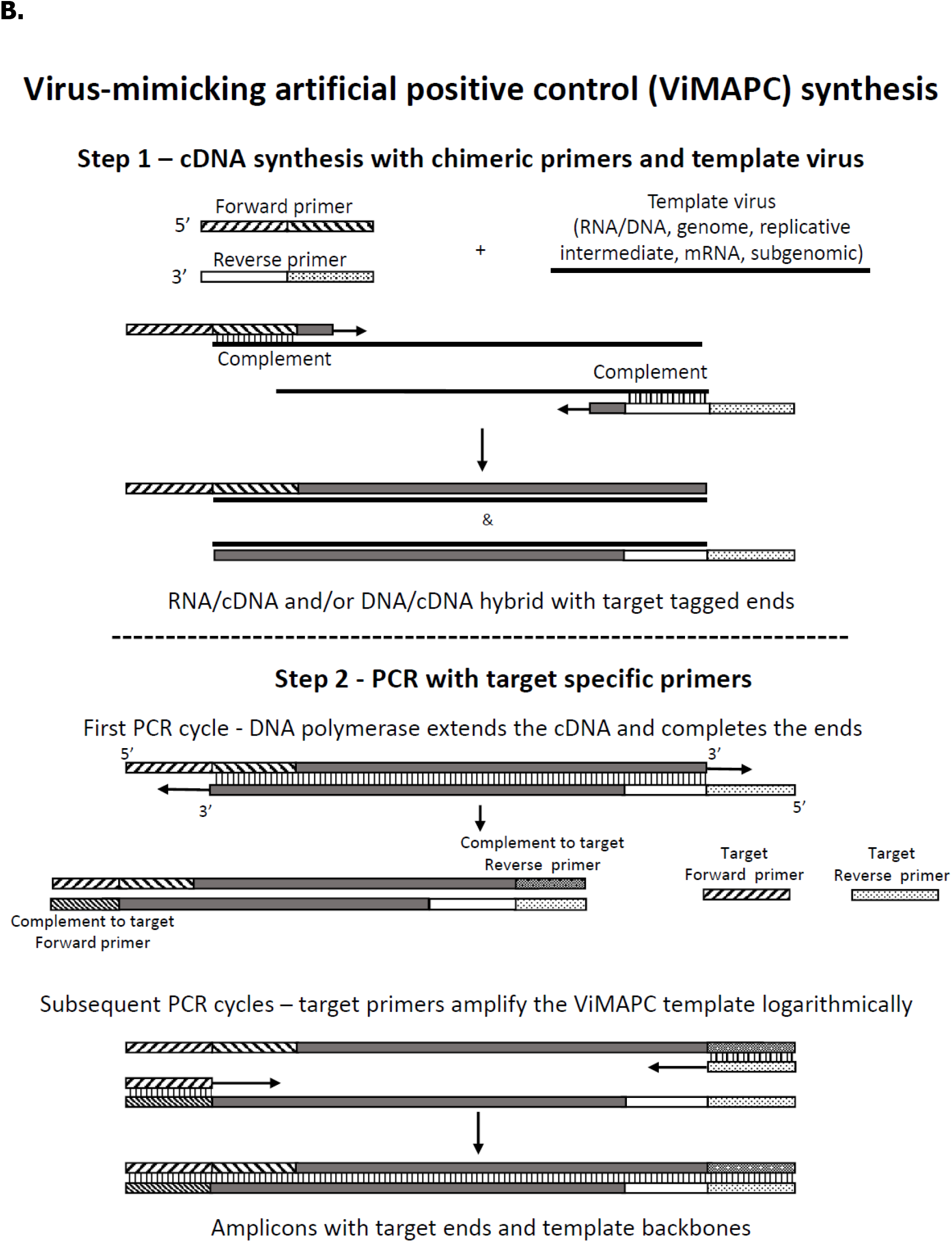
Overview and concept of primers used for the development of virus-mimicking artificial positive control (ViMAPC). A: Overview of ViMAPC development and their utilization in diagnostics; a: virus templates available in different genome composition/types: cartoon depiction of the primers used in reverse transcription; c: pictorial representation of the ViMAPC developed; d: expected outcome of a target virus PCR with tissue naturally infected with the target virus and the ViMAPC developed for the same. B: Development of ViMAPCs. Step 1: Reverse transcription (RT) using template virus (RNA/DNA genome) and primers (target + template) resultant into RNA/cDNA and/or DNA/cDNA with overhang target virus ends. Step 2: Polymerase chain reaction (PCR) of the resultant RNA/DNA and/or DNA/cDNA material. In the first PCR cycle the cDNA stands denature and the single stranded cDNA molecules reanneal forming a double stranded cDNA molecule with target primer overhangs at each end. The DNA polymerase fills in the ends. In subsequent cycles the primers for the target virus bind to the completed ViMAPC template and amplify logarithmically.

Based on the fact that the reverse transcriptase is an RNA and DNA dependent DNA polymerase (Tzanetakis et al., 2005), the template could be either RNA and/or DNA. During RT, the 3’ portion of the primer, corresponding to the template, binds to the nucleic acids (genomic, replication intermediates or mRNA). The cDNA is extended resulting in RNA-cDNA (RNA viruses) or DNA/cDNA + RNA/cDNA hybrids (DNA viruses) with the 5’ end of the primer, corresponding to the target, forming an overhang at the 5’ of the newly created strand (Fig.1B).

#### The first PCR cycle is critical in the process

During denaturation the RT-generated hybrid strands separate. At the annealing step the cDNA obtained from the +/-RNA and/or DNA strands form dsDNA whilst the DNA (Taq) polymerase fill in the 5’ overhangs during extension (Fig. 1B). The repaired molecule then amplifies exponentially using the target primers in subsequent PCR cycles. To examine whether the RT primers affected amplification, and to test the applicability and effectiveness of the technology RNA and DNA viruses were used in proof-of-concept experiments (Table 1; Fig. 2).

**Table 1.**
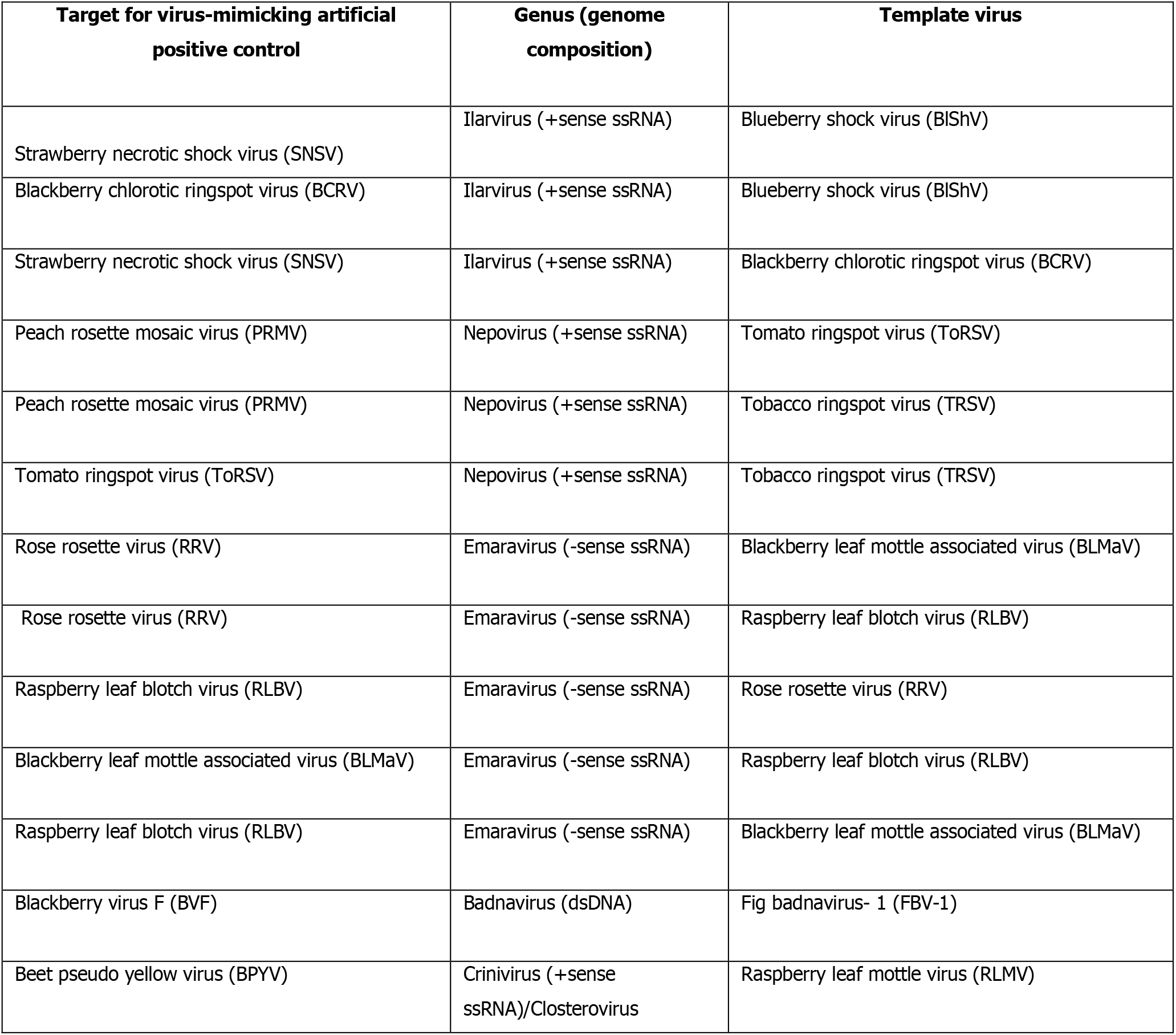
List of targets and their corresponding virus-infected templates used for preparation of virus-mimicking artificial positive controls (ViMAPCs). The genus and genome composition for both target and template are provided. Acronyms: dsDNA: double stranded DNA; -ssRNA: negative sense single stranded RNA; +ssRNA: positive sense single stranded RNA.

**Figure 2.**
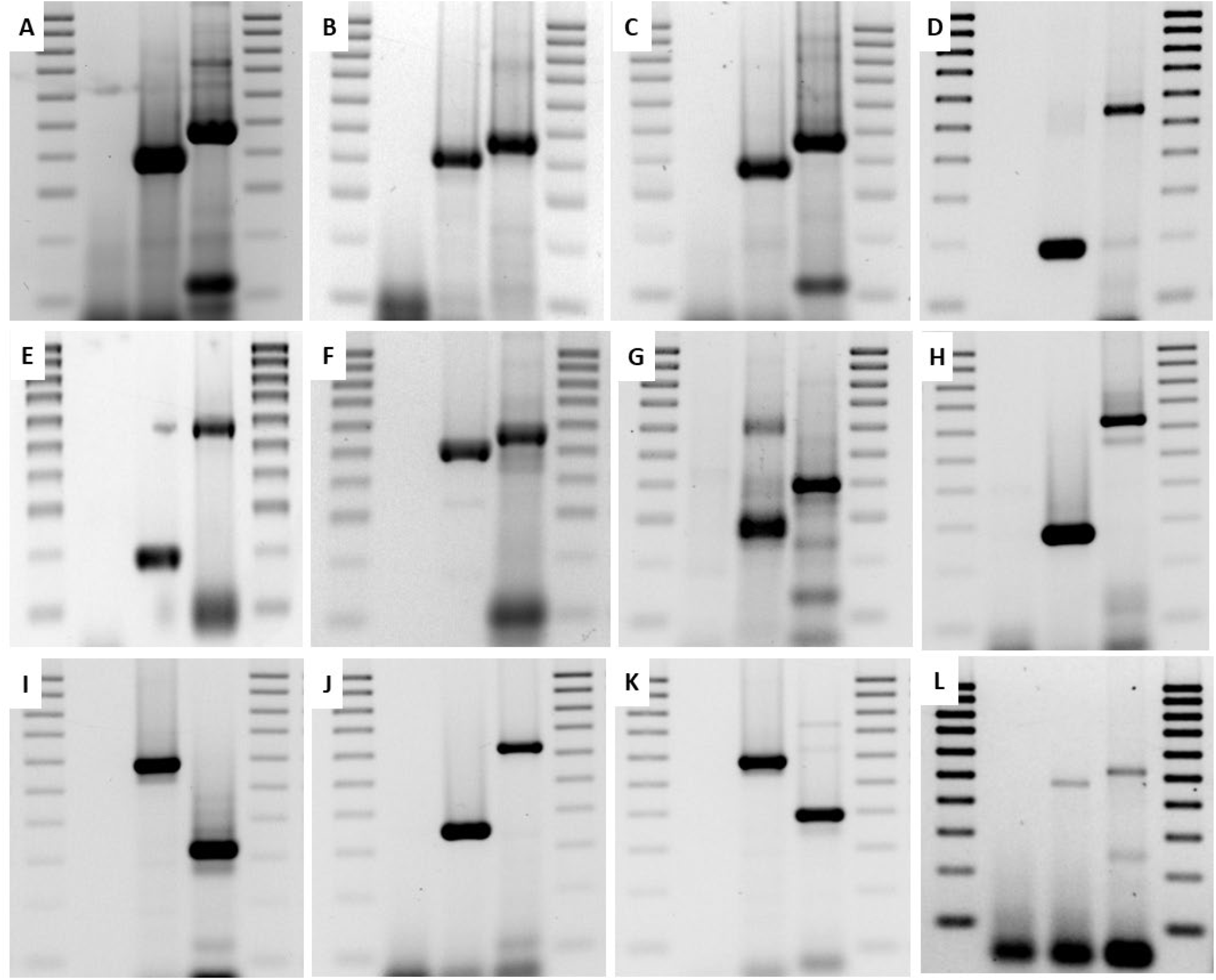
Verification of virus-mimicking artificial positive controls (ViMAPCs) using either RNA or DNA as template (please refer to Table 1 for taxonomic placement of the viruses). Each gel presents the amplicon of a specific target virus, with a plant negative, plant positive and the developed ViMAPC using as template material infected with a virus belonging to the same genus. Lane 1 = 100 bp DNA marker; Lane 2 = Plant negative for target virus; Lane 3= Plant positive for target virus; Lane 4 = ViMAPC and Lane 5 = 100 bp DNA marker. **A**. strawberry necrotic shock virus (SNSV) using blueberry shock virus (BlShV) as template; **B**. blackberry chlorotic ringspot virus (BCRV) using BlShV as template; **C**. SNSV using BCRV as template; **D**. peach rosette mosaic virus (PRMV) using tomato ringspot virus (ToRSV) as template; **E**. PRMV using tobacco ringspot virus (TRSV) as template; **F**. ToRSV using TRSV as template; **G**. rose rosette virus (RRV) using blackberry leaf mottle associated virus (BLMaV) as template; **H**. RRV using raspberry leaf blotch virus (RLBV) as template; **I**. RLBV using RRV as template; **J**. BLMaV developed using RLBV as template **K**. RLBV using BLMaV as template and **L**. blackberry virus-F (BVF) using fig badnavirus-1 (FBV-1) as template.

### Reverse transcription PCR

RT reaction mixture (5X RT Buffer, dNTPs (0.4 mM), primers (400nM, Supplemental table 2), template and molecular biology grade water to 25 μl) were incubated at 70°C for 5 min. The reaction tube was placed on ice for 5 min, followed by the addition of Maxima® reverse transcriptase (50U) and RNAse inhibitors (6U) (ThermoFisher Scientific). The reaction was then incubated at 50°C for 60 min, followed by heat inactivation at 85°C for 5 min. Plant material other than that used for development of ViMAPCs were reverse transcribed with random primers following the protocols used for diagnostics in our laboratory (Thekke-Veetil et al., 2016).

PCR reactions consisted of 10X Taq buffer, primers (400nM), dNTPs (0.4 mM), RT product (2% vol/vol), Taq DNA polymerase (0.5U, Genescript), and molecular biology grade water to 25μL. For each reaction, the previously published parameters for the target virus were applied (Supplemental table 1). Amplicons were separated using 1X SB buffer on a 1.5% agarose gel, followed by post-staining with 1X GelRed, visualized on UV transilluminator and captured using Gel Doc™ (BioRad).

### Application

For conceptual verification, PCR was conducted for each virus on target-infected (true positive) and target-negative material (true negative) along with the ViMAPC. We tested ViMAPCs in cross-host combinations; considered a suboptimal scenario for a ViMAPC effectiveness. As a mock for a standard diagnostic test *Rubus* and *Fragaria* samples from the NCGR collection were tested for the presence of strawberry necrotic shock virus (SNSV) as described (Thekke-Veetil et al., 2016; Fig. 3). The ViMAPC for SNSV (target) was developed using blueberry shock virus (BlShV) (template) infecting *Vaccinium* and the primer sequences described in Thekke-Veetil et al., (2016) and Villamor et al. (2022) for SNSV and BlShV detection, respectively.

**Figure 3.**
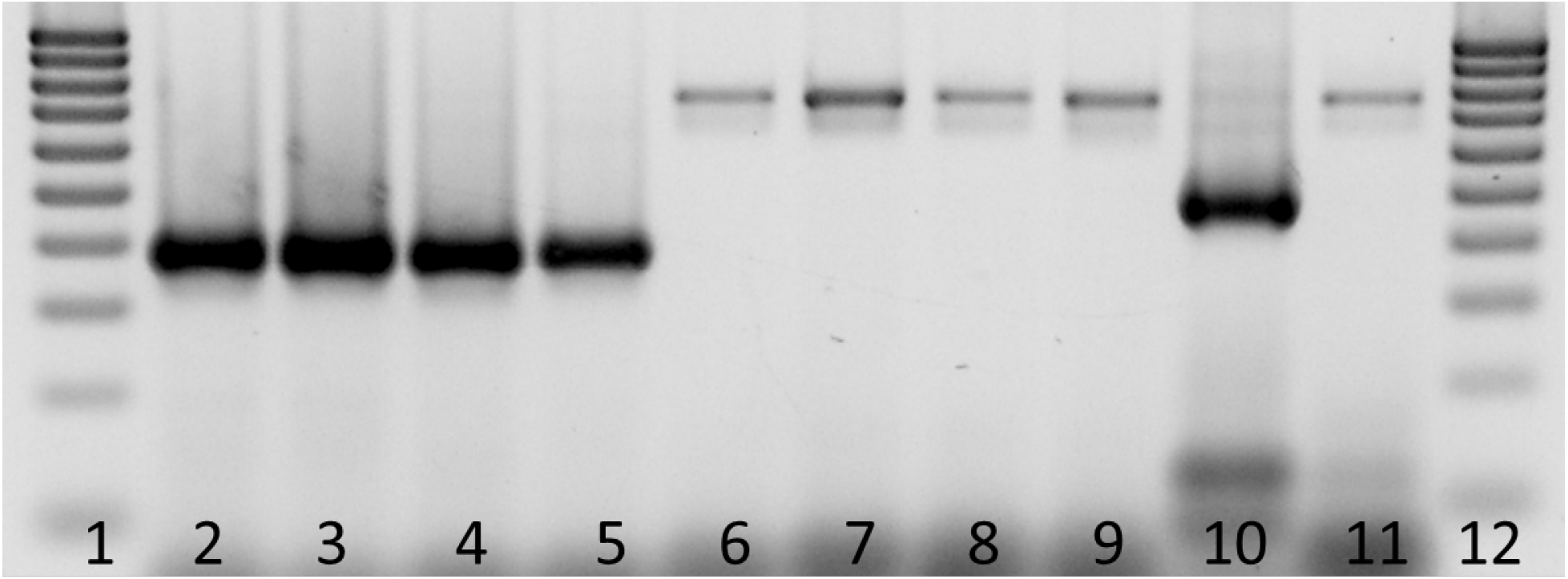
Application and detection of strawberry necrotic shock virus (SNSV) virus-mimicking artificial positive control (ViMAPC) developed using blueberry shock (BlShV)-infected material as template. Material infected with SNSV (*Rubus:* PI 691930 and PI 691943, *Fragaria* : PI 691784; PI 691800) and previously tested negative for the virus (*Rubus* : PI 691933 and PI 691936; *Fragaria* : PI 691758 and PI 691766) were tested alongside the ViMAPC (Lane 10) (supplementary table 3). All lanes show expected band sizes (supplementary table 1 and 2) (Thekke-Veetil et al., 2016). Lane 1 and Lane 12 represented 100 bp DNA marker.

## RESULTS AND DISCUSSION

### Concept

Twelve homologous (template/target belong to the same virus genus) and one heterologous (template/target belong to different genera) ViMAPCs were developed and applied successfully on both RNA and DNA viruses. It is important to note that utilizing RT enhances the detection sensitivity for DNA viruses because of the relative abundance of virus RNA when compared to genomic DNA (Laney et al., 2012; Shahid et al., 2017).

The effect of the RT primers in amplification was monitored by comparing the same virus as a target of a diagnostics PCR and as the template of a ViMAPC. Overall, results indicate that the primers do not have a major effect in the process (Fig. 3) with ViMAPC and corresponding true positive yielding amplicons of similar intensity; yet there were exceptions.

The use of published primer sequences allows for the rapid design of the RT primers and builds off already validated tests. Therefore, when a ViMAPC is required rapidly, such as during a suspected incursion, the use of published primers accelerates the process. However, it should be noted that the RT primers may lead to RT kinetics that are suboptimal (because of the secondary structure of the RT primer that may not be fully melted at the RT denaturation step or misannealing because of the high melting primer temperature) which may result in differences in intensity between true positive and ViMAPC. This could be addressed either by modifications in the RT protocol or by designing primers that bind optimally to their template during the RT (tobacco ringspot virus, Supplemental Table 1). When ViMAPC results in suboptimal amplification diagnosticians need to decide based on their needs and context of the experiment. For example, peach rosette mosaic virus-infected tissue is extremely difficult to obtain because of the limited distribution of the virus, yet by using the described strategy a ViMAPC was consistently amplified even though not to the intensity obtained from a natural infection (Fig. 2).

Another important point is that template and target viruses need to be closely related to obtain amplicons which optimally mimic virus infection. When using related yet phylogenetically paraphyletic viruses (e.g., different genera in the same family) results can be are suboptimal (Supplemental Fig. 1).

### Application

The ViMAPC concept was applied in a real-life scenario for the detection of SNSV. *Rubus* and *Fragaria* samples were tested for SNSV whereas the ViMAPC was developed using BlShV-infected *Vaccinium* as template. Natural SNSV infection yields a 372bp amplicon whereas the region between the priming site of the template (BlShV) is 450bp.

The additional bases added in the ViMAPC (24+25 bases for SNSV forward/reverse primers) result to a 499bp product making differentiation between natural infection and ViMAPC contamination easily identifiable by observing amplicon migration on an agarose gel (Fig. 3).

We have provided evidence that the approach presented in this communication mimics virus infection and also eliminates the possibility of misinterpreting contamination as natural infection. ViMAPCs provide the only known approach to date where an artificial control mimics natural virus infection. In addition, this technology does not necessitate any additional skills from the diagnostician, is cost effective, solely requiring the synthesis of the two oligonucleotide primers, and can be applied in a fraction of the time compared to the alternatives making a tool that can be used in all cases, including those where infected material is difficult to obtain.

ViMAPC will accelerate the speed for diagnostics. As a proof-of-concept we used plant virus-infected material that was readily available to this group. This technology is readily applicable to all organisms as we have shown it can be used on DNA and RNA templates as long as there is a related organism available to be used as the template. The mimicking aspect of the technique and the fact that it can be easily and quickly deployed suggests it can be very useful when working with quarantine pathogens, or in cases where maintaining a large collection of positive controls is impractical.

## Author contribution

The design of the study was done by I.E.T. The experiments were performed by S.S. and data analyses were done by S.S. D.S and I.E.T. The manuscript was drafted, edited and revised by all authors.

---

**Supplemental Table 1.**
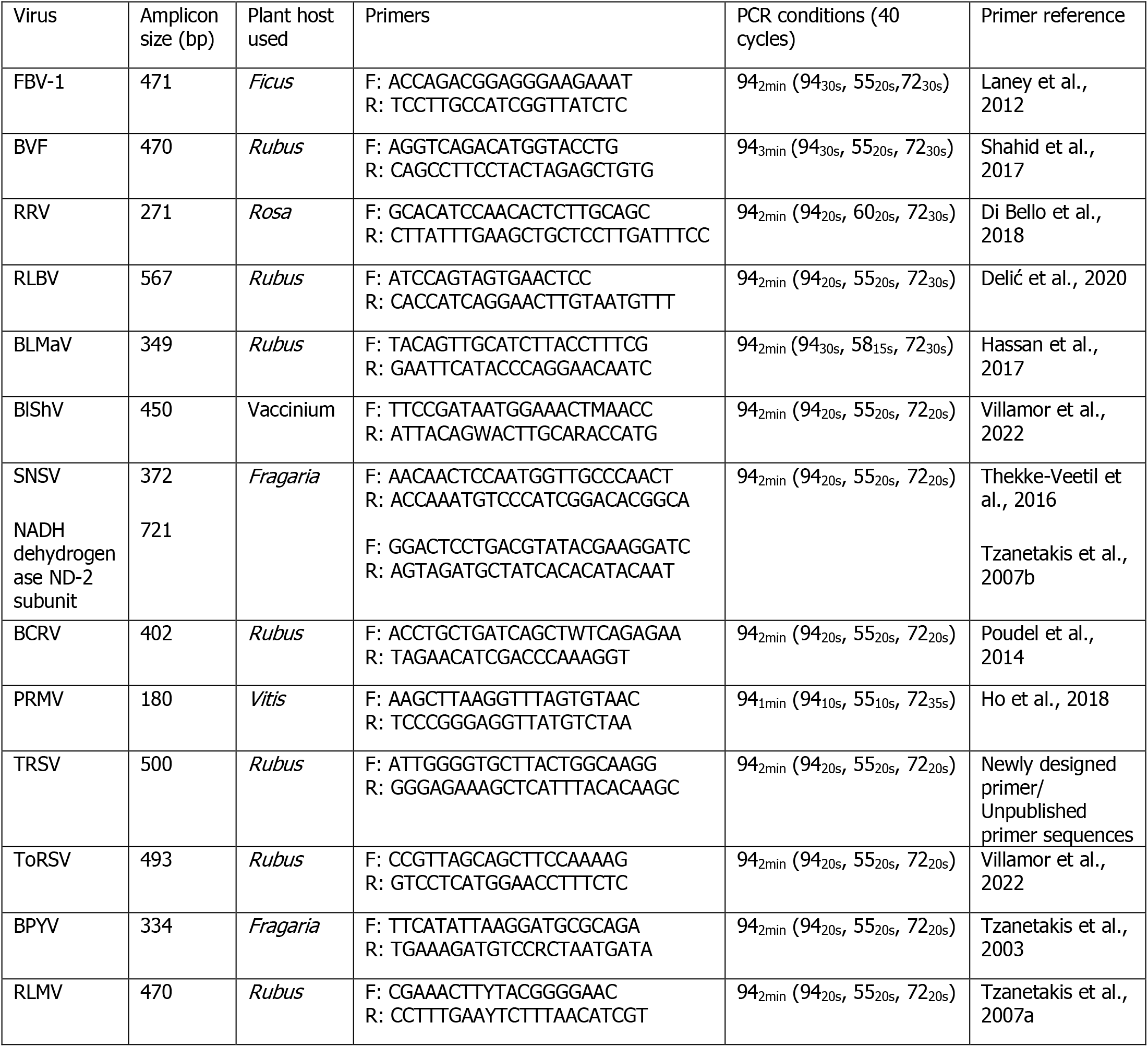
List of detection primers with their optimized PCR protocols for virus detection. BCRV: blackberry chlorotic ringspot virus; BLMaV: blackberry leaf mottle associated virus; BlShV: blueberry shock virus; BPYV: beet pseudo yellows virus; BVF: blackberry virus F; FBV-1: fig badnavirus-1; PRMV: peach rosette mosaic virus: RRV: rose rosette virus; RLBV: raspberry leaf blotch virus; RLMV: raspberry leaf mottle virus SNSV: strawberry necrotic shock virus; ToRSV: tomato ringspot virus and TRSV: tobacco ringspot virus.

**Supplemental Table 2.**
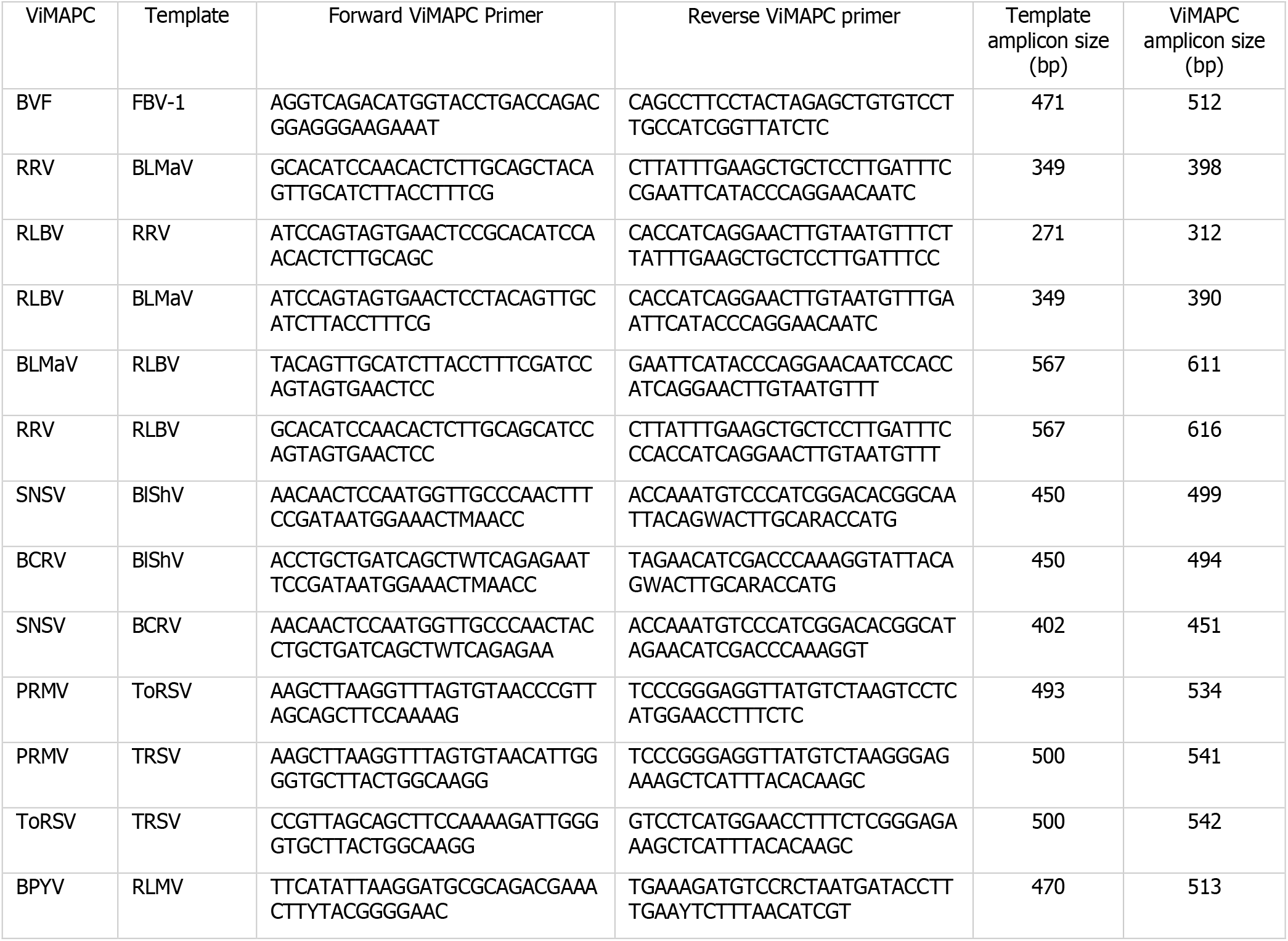
Reverse transcription primers for the development of virus-mimicking artificial positive controls (ViMAPC) and expected amplicon size. Acronyms: BCRV: blackberry chlorotic ringspot virus; BLMaV: blackberry leaf mottle associated virus; BlShV: blueberry shock virus; BPYV: beet pseudo yellows virus; BVF: blackberry virus F; FBV-1: fig badnavirus-1; PRMV: peach rosette mosaic virus: RRV: rose rosette virus; RLBV: raspberry leaf blotch virus; RLMV: raspberry leaf mottle virus SNSV: strawberry necrotic shock virus; ToRSV: tomato ringspot virus and TRSV: tobacco ringspot virus

**Supplemental Figure 1.**
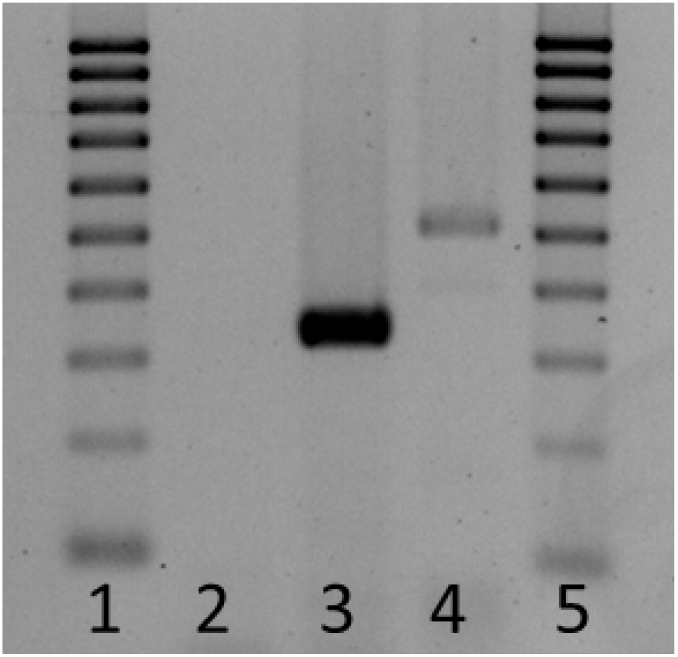
Virus-mimicking artificial positive control (ViMAPC) for beet pseudo yellows virus (BPYV, genus *Crinivirus, family Closteroviridae*) developed using raspberry leaf mottle virus (RLMV, genus *Closterovirus, family Closteroviridae*) as template. Lanes 1 and 5: 100 bp DNA marker; lane 2: negative control; lane 3: positive control, 334 nucleotides; lane 4: BPYV ViMAPC, 513 nucleotides.

